# Combination of *in vivo* phage therapy data with *in silico* model highlights key parameters for treatment efficacy

**DOI:** 10.1101/2021.03.04.433924

**Authors:** Raphaëlle Delattre, Jérémy Seurat, Feyrouz Haddad, Thu-Thuy Nguyen, Baptiste Gaborieau, Rokhaya Kane, Nicolas Dufour, Jean-Damien Ricard, Jérémie Guedj, et Laurent Debarbieux

## Abstract

The clinical (re)development of phage therapy to treat antibiotic resistant infections requires grasping specific biological properties of bacteriophages (phages) as antibacterial. However, identification of optimal dosing regimens is hampered by the poor understanding of phage-bacteria interactions *in vivo.* Here we developed a general strategy coupling *in vitro* and *in vivo* experiments with a mathematical model to characterize the interplay between phage and bacterial dynamics during pneumonia induced by a pathogenic strain of *Escherichia coli.* The model estimates some key parameters for phage therapeutic efficacy, in particular the impact of dose and route of administration on phage dynamics and the synergism of phage and the innate immune response on the bacterial clearance rate. Simulations predict a low impact of the intrinsic phage characteristics in agreement with the current semi-empirical choices of phages for compassionate treatments. Model-based approaches will foster the deployment of future phage therapy clinical trials.

## Introduction

The pronounced slowdown in the discovery of new antibiotics to treat bacterial infections caused by multidrug resistant (MDR) pathogens has revived the interest for bacteriophages (phages), viruses infecting bacteria (Gordillo Altamirano and Barr, 2019; Kortright et al., 2019). Pre-clinical experiments and several clinical cases reports have demonstrated the potential of phages to successfully treat infections caused by MDR bacteria (Aslam et al., 2020; Corbellino et al., 2020; Jennes et al., 2017; Melo et al., 2020; Schooley et al., 2017). These success are nonetheless balanced by clinical studies that did not confirm these positive results (Jault et al., 2019; Leitner et al., 2020; Sarker et al., 2016), showing that translation from experimental results and promising individual treatments to clinical trials remains challenging.

A major milestone for a widespread use in human populations is the determination of the optimal dose, route of administration and treatment duration. This is a particularly complex endeavor for phages, as standard assessments of clinical pharmacology used to determine the processes of Administration, Distribution, Metabolism and Excretion (ADME) of drugs are not adapted to phages. Indeed, phages massively replicate on their target bacteria with kinetic parameters governing this viral amplification (receptor recognition, genome amplification, particles assembly and bacterial lysis) being specific to each phage-bacteria pair (Dion et al., 2020). Furthermore, phages are naturally present in human-associated microbiotas and their rate and route of elimination do not follow usual metabolic pathways through kidney or liver, as for xenobiotic products (Dabrowska and Abedon, 2019).

Here we coupled *in vitro* and *in vivo* experiments with mathematical modeling to approach the key parameters of phage-bacteria-host interactions in order to guide treatment strategies. This approach has been successful in drug development for various chronic (HIV, HCV) and acute (Ebola, Zika, Influenza) viral infections (Friberg and Guedj, 2020; Perelson and Guedj, 2015). Unlike previous models that focused on specific aspects of phage-bacteria, phage-host immune system or bacteria-host immune system interactions (Cairns et al., 2009; Roach et al., 2017; Smith et al., 2011; Weld et al., 2004), we aimed to design a synthetic, semi-mechanistic model that could be used to understand the tripartite phage-bacteria-host interactions *in vivo*. We purposely designed a series of specific experiments (Figure 1) to estimate the central parameters from a previously validated experimental mice model of pulmonary infection using the bioluminescent *Escherichia coli* pathogenic strain 536 and the virulent phage 536_P1 (Dufour et al., 2015). Dynamic and direct microbiological data were collected from groups of animals that were either (bacterial) uninfected and (phage) treated, infected and untreated, or infected and treated, using different inoculum doses and routes of administration. These data led us to develop an original mathematical model recapitulating the tripartite interactions over time. This model identified key parameters affecting bacterial kinetics *in vivo*. It was then exploited to test scenarios including variations of phage intrinsic properties. The use of such model will rationalize the phage choice, the dose and route of administration in order to optimize the efficacy of phage therapy.

**Figure 1.**
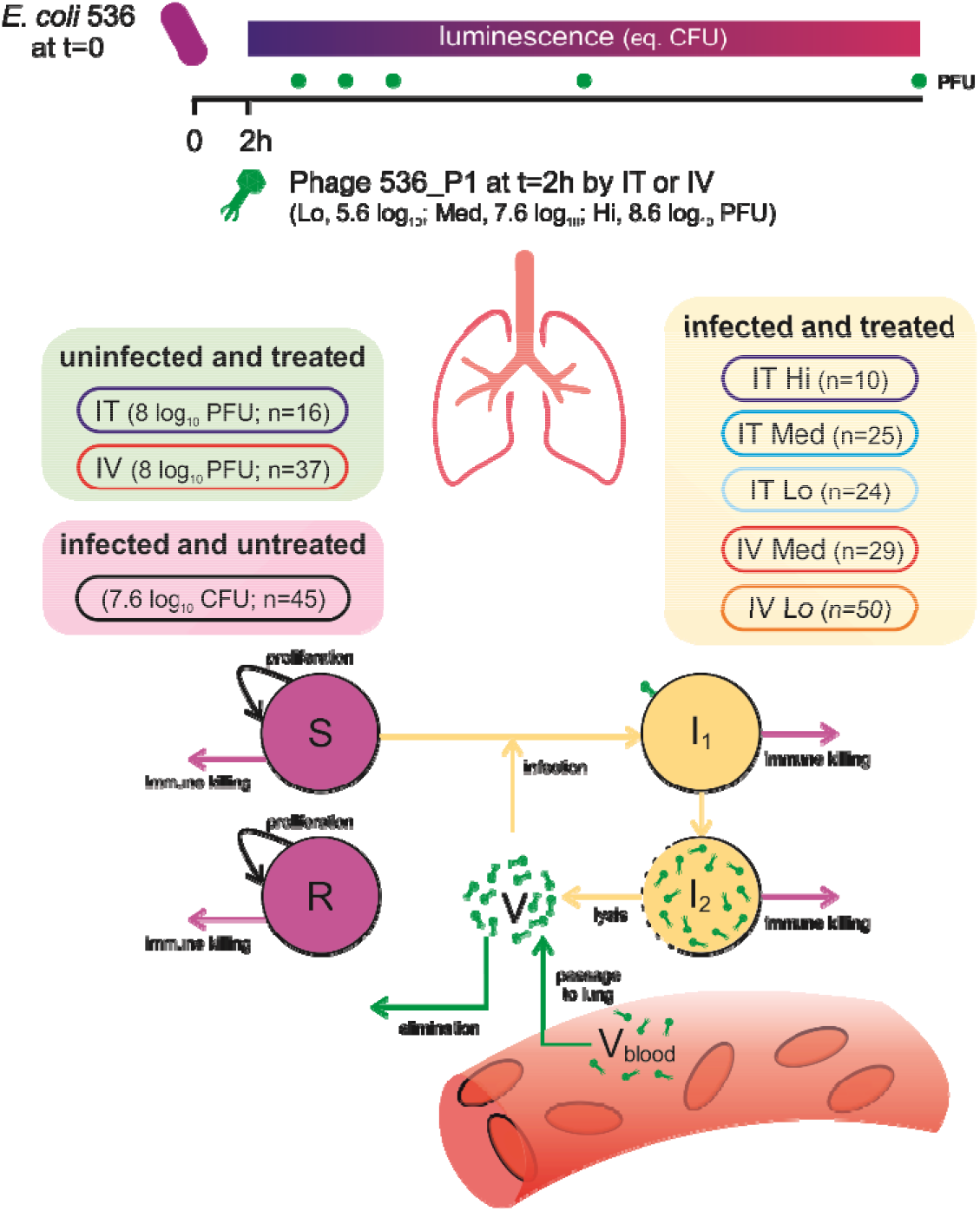
Schematic representation of the experimental design of the study. *In vivo* experiments were conducted in mice that were either i) uninfected and treated (green box, with phages administered either intratracheally, IT, or intravenously, IV); ii) infected and untreated (magenta box); iii) infected and IT- or IV-treated at various doses (yellow box). The intensity of light emitted by *E. coli* strain 536-lux was recorded along all experiments and translated to bacterial lung density, while phage 536_P1 density was measured only once (upon sacrifice) in each animal at specific time points. A mathematical compartmental model was developed to fit to these data, with S, R, I1 and I2 representing the susceptible, refractory, infected non-productive, productively infected bacteria, respectively, and V and Vblood the phage concentration in lungs and blood, respectively.

## Results

### Biodistribution of phage in uninfected mice

We first studied the biodistribution of phage 536_P1 in uninfected mice following a single administration of 8 log_10_ PFU by intratracheal (IT) or intravenous (IV) route (Figure 2). After IT administration, phage concentrations in the lungs varied over time only within 1-log with observed mean values between 7.0 and 8.0 log_10_ PFU/g. By contrast, after IV administration, the observed peak concentration was reached four hours post-administration (5.1 [4.7, 5.5] log_10_ PFU/g), which was over 2-log below the observed concentration after IT administration (7.2 ± [6.8, 7.8] log_10_ PFU/g). Therefore, after IV administration only a fraction *F* of the phage, estimated to 0.02%, ultimately reached the lung compartment, leading to exposure approximately 5000 times lower than after IT administration and an estimation of the absorption rate into the lung compartment, *k_a_,* of 0.3 h^−1^. (Eq. 1-2). The kinetics of phage elimination was also different with an estimated half-life of 12 and 3 hours after IT and IV administration, respectively (see *k_e_*(IT) and *k_e_*(*V*) values in Table 1).

**Figure 2.**
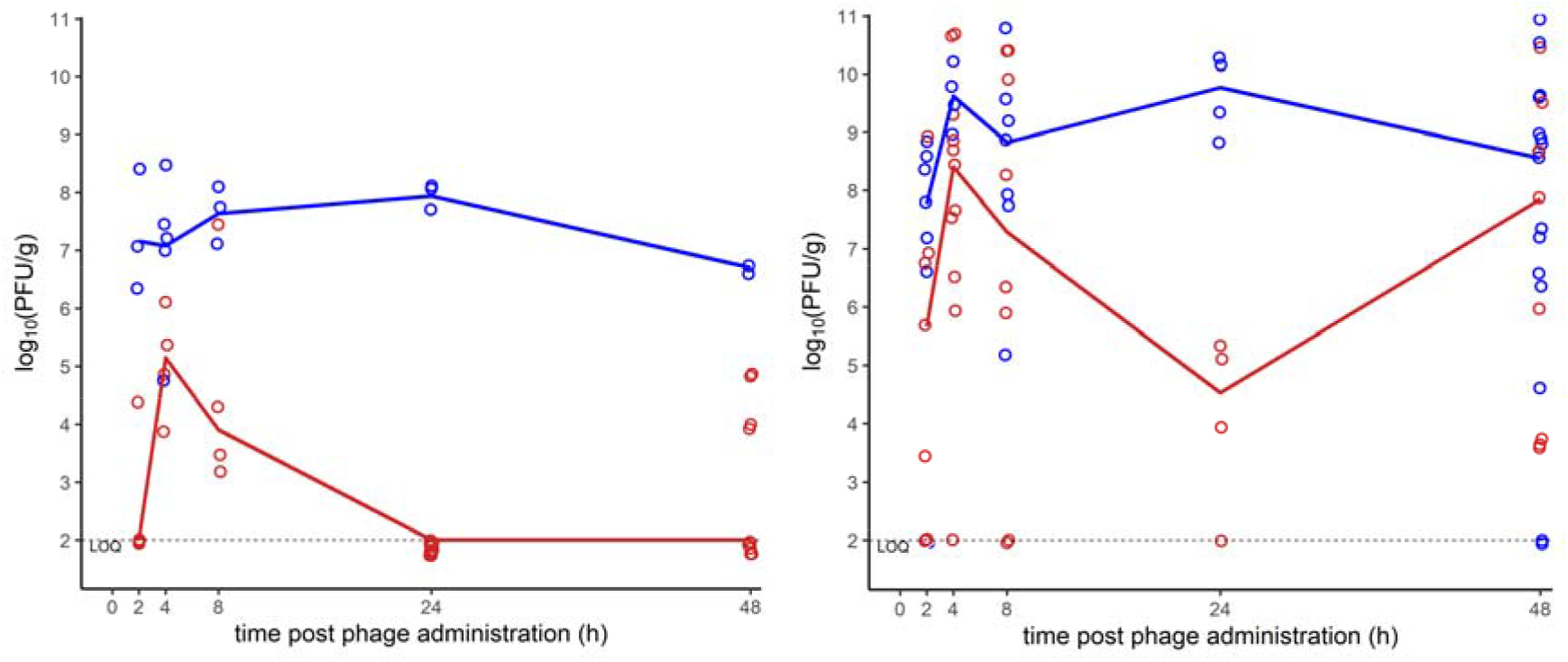
Distribution of phage 536_P1 in lungs of treated uninfected and treated infected mice. Lungs of uninfected (left, N=53) and *E. coli* infected (right, N=138) mice were homogenized at 2, 4, 8, 24 or 48h post-administration of a single dose of phage 536_P1 (8 log_10_ PFU in uninfected mice and 5.6, 7.6, or 8.6 log_10_ PFU in infected mice), either by intratracheal (blue) or intravenous (red) route. Lines represent the median phage concentrations at each time point. The dashed line represents the limit of quantification (LOQ=2 log_10_ PFU/g).

**Table 1.** Parameter estimates of the final phage/bacteria dynamics model in mice

| Parameter | Name | Unit | Fixed/Estimated from | Fixed effect (rse, %) | Sd of random effect (rse, %) |
| --- | --- | --- | --- | --- | --- |
| Bacterial growth rate | $\alpha$ | $\text{h}^{-1}$ | Untreated infected | 0.21 (-) | 0 (-) |
| Initial number of bacteria | $B_0$ | $\log_{10} \text{CFU g}^{-1}$ | Untreated infected | 7.1 (1) | 0.14 (6) |
| Bacterial load plateau | $B_{max}$ | $\log_{10} \text{CFU g}^{-1}$ | Untreated infected | 10.4 (-) | 0 (-) |
| Time for immunity activation | $t_I$ | h | Untreated infected | 3.0 (-) | 0 (-) |
| Maximal immunity effect | $\delta_M$ | $\text{CFU g}^{-1} \text{h}^{-1}$ | Untreated infected | $6.7 \cdot 10^6$ (-) | 0.05 (-) |
| Bacterial load for 50% of $\delta$ | $B_{50}$ | $\text{CFU g}^{-1}$ | Untreated infected | $2.3 \cdot 10^7$ (-) | 0 (-) |
| Infectivity rate | $\beta$ | $\log_{10} \text{g CFU}^{-1} \text{h}^{-1}$ | Treated infected | 6.9 (7) | 0.11 (43) |
| Transition rate | $e$ | $\text{h}^{-1}$ | *in vitro* | 1.2 (-) | 0 (-) |
| Lysis rate | $\delta_V$ | $\text{h}^{-1}$ | Treated infected | 0.09 (25) | 0 (-) |
| Burst size | $b$ | $\text{PFU CFU}^{-1}$ | *in vitro* | 570 (-) | 0 (-) |
| Relative bioavailability (IV/IT) | $F$ | - | Treated uninfected | 0.0002 (-) | 0 (-) |
| Phage absorption rate | $k_a$ | $\text{h}^{-1}$ | Treated uninfected | 3 (-) | 0 (-) |
| Phage elimination rate after IT | $k_e(IT)$ | $\text{h}^{-1}$ | Treated uninfected | 0.058 (-) | 0 (-) |
| Phage elimination rate after IV | $k_e(IV)$ | $\text{h}^{-1}$ | Treated uninfected | 0.23 (-) | 0 (-) |
| Phage distribution weight | $W$ | g | Treated uninfected | 1 (-) | 0 (-) |
| Initial refractory proportion | $f$ | - | Treated infected | 0.105 (45) | 2.8 (19) |
| Weibull shape | $\kappa$ | - | Treated infected | 1.8 (11) | 0 (-) |
| Weibull scale | $\lambda$ | $\text{h}^{-1}$ | Treated infected | $2.4 \cdot 10^4$ (98) | 0 (-) |
| Impact of bacterial kinetics on death | $\gamma$ | $\text{g CFU}^{-1}$ | Treated infected | 1.44 (13) | 0 (-) |
| Additive error on bacterial load | $\sigma_B$ | $\log_{10} \text{CFU g}^{-1}$ | Treated infected | - | 0.47 (4) |
| Additive error on phage concentration | $\sigma_V$ | $\log_{10} \text{PFU g}^{-1}$ | Treated infected | - | 2.6 (11) |

Sd: standard deviation, rse: relative standard error

### Bacterial kinetics in infected-untreated mice

IT administration of the *E. coli* strain 536 led to a large variability in the lung bacterial load of untreated animals, as measured by emitted luminescence and CFUeq/g (see methods), with observed values ranging between 5.0 and 9.9 log_10_ CFUeq/g at two hours post infection (Figure 3). This heterogeneity in the effective bacterial inoculum mimics the diversity of bacterial loads in human patients with pneumonia (Jiang et al., 2014), and allowed us to consider each mouse as an individual when developing the mathematical model (see below and Eq. 3). This heterogeneity also informed on the threshold of bacterial density above which the immune response of the host cannot control the infection. Indeed, when the bacterial load was greater than 6.8 log_10_ CFUeq/g at two hours post-infection, the bacterial density increased continuously, while, below this threshold, we observed a rapid reduction of the bacterial density caused by the immune response to the infection (Figure 3).

**Figure 3.**
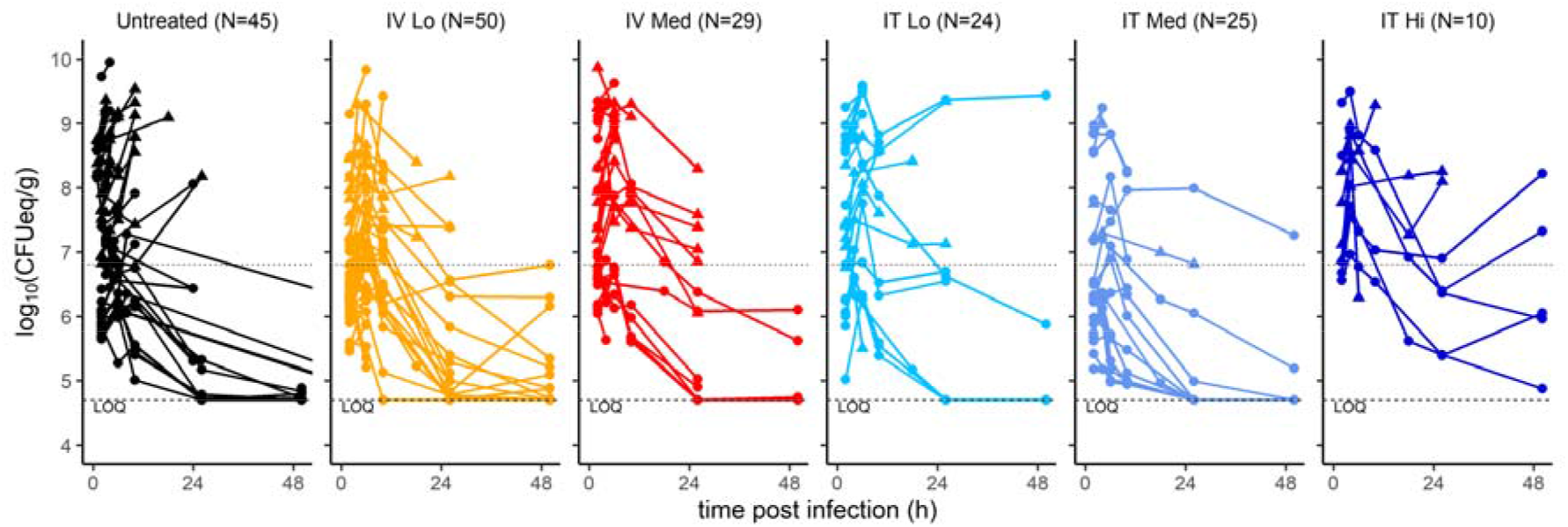
Kinetics of bacterial densities in untreated and phage-treated animals. Bioluminescence data from infected animals were stratified by treatment groups: untreated (black), treated with phage 536_P1 administered by IV at a low dose (5.6 log_10_ PFU, yellow) or medium dose (7.6 log_10_ PFU, red), or by IT at either low dose (cyan), or medium dose (blue) or high dose (8.6 log_10_ PFU, dark blue). The dotted horizontal line at 6.8 log_10_ CFUeq/g corresponds to the threshold value of bacterial load at two hours post-infection below which all animals controlled the infection. The dashed line represents the limit of quantification (LOQ=4.7 log_10_ CFUeq/g). Circles correspond to live animals until the time of sacrifice while triangles correspond to dead animals before the scheduled time of sacrifice.

### Bacterial and phage kinetics in infected treated mice

Next, using the same infection procedure as above, mice (N=138) were treated two hours post-infection with various doses of phage 536_P1 (low dose: 5.6; medium dose: 6.6; high dose: 7.6 log_10_ PFU), administered either by IT or IV route.

Phages rapidly replicated and their density in the lungs reached values 2 to 3-log_10_ higher than in uninfected animals, showing that they encountered bacteria and efficiently replicated (Figure 2). At four hours post-phage administration (six hours post-bacterial infection), the median titer of phage in the lungs of IT-treated mice (N=59) was 9.6 [9.3, 9.9] log_10_ PFU/g, as compared to 7.1 [7.1, 7.4] log_10_ PFU/g in uninfected IT-treated mice (N=16). A similar difference was also observed in IV-treated animals (N=79), with phages concentration of 8.4 [7.0, 9.1] log_10_ PFU/g as compared to 5.1 [4.7, 5.5] log_10_ PFU/g in uninfected IV-treated animals (N=37). A dose effect was noticed in IV-treated animals, with a median concentration of 7.6 [6.4, 8.5] log_10_ PFU/g in the lungs of mice receiving a low dose (N=50), as compared to 10.6 [10.0,10.6] log_10_ PFU/g in mice treated with a medium dose (N=29).

The initial effective median bacterial inoculum was similar in each of the three groups with levels of 6.9 [6.1, 8.0], 7.2 [6.3, 8.5] and 6.7 [6.3, 7.8] log_10_ CFUeq/g in the lungs of untreated, IT- and IV-treated and animals, respectively (Figure 3). Afterwards, two hours post-phage administration, the bacterial loads tended to be lower in treated (N=138) compared to untreated (N=45) animals, with levels of 7.5 [6.6, 8.8] log_10_ CFUeq/g and 8.4 [7.6, 8.8] log_10_ CFUeq/g, respectively (p=0.17, Wilcoxon-Mann-Whitney test). However, at this time point there was no effect of the route of administration, with values of 7.5 [6.5, 8.8] log_10_ CFUeq/g and 7.2 [6.6, 8.5] log_10_ CFUeq/g in IT and IV-treated groups, respectively (p=0.27 and p=0.14 to untreated, respectively, and p=0.75 when comparing IT and IV-treated groups, Wilcoxon-Mann-Whitney test).

The mortality rates at 48 hours were similar in all groups (respectively 42%, 48% and 52% in IT, IV and untreated animals) (Figure 4). Mortality rate was not dependent on the dose of administered phages (at 48 hours equal to 51%, 25% and 50% in animals receiving a low, medium or high dose of phages, respectively; p=0.92, 0.14 and 0.68 to untreated, logrank test), (Figure S4). However, it is important to note that in the subgroup of animals with a high effective bacterial inoculum (>6.8 log_10_ CFUeq/g), treatment tended to increase the survival rate (Figure 4, p=0.07 in IT and p=0.21 in IV, as compared to untreated, respectively, logrank test), whereas all animals with a low effective inoculum (<6.8 log_10_ CFUeq/g) survived at 48h, regardless of treatment (Figure 4).

**Figure 4.**
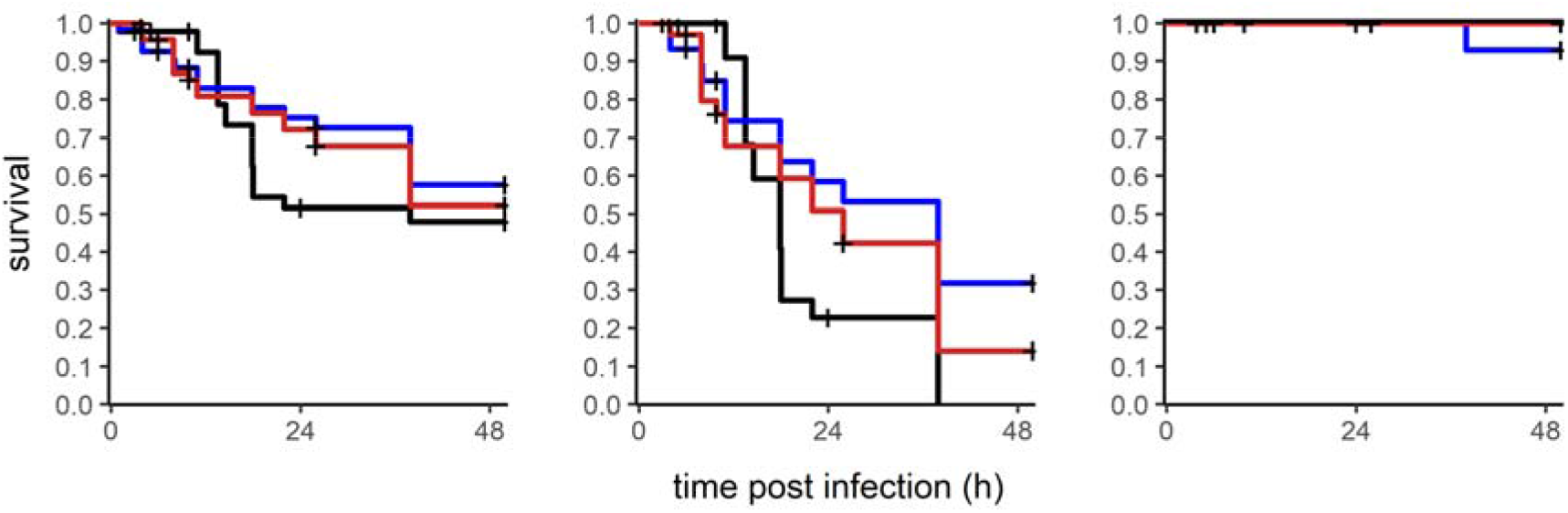
Survival of infected and treated animals in function of the pulmonary bacterial load at two hours post-infection. The survival curves were obtained from animals that were infected untreated (black, N=45), infected treated by IT route (blue, N=59) or infected treated by IV route (red, N=79). The left panel displays all infected animals (N=183). The middle panel includes only animals (N=78) with a pulmonary bacterial load at two hours post-infection greater than 6.8 log_10_ CFUeq/g. The right panel includes only animals (N=99) with a pulmonary bacterial load at two hours post-infection lower than 6.8 log_10_ CFUeq/g. Crosses represent sacrifice time. Of note, 6 animals sacrificed at two hours post infection were not represented in both middle and right panel.

### Unraveling bacteria-phage interaction using a mathematical model

Next, we used these data in both infected and uninfected animals together with parameters determined *in vitro* (see methods) to build a mathematical model of phage-bacteria-host interaction (Eq. 4-10, Figure 1). Fitting the model to bacterial load data, we first determined that a model accounting for the control of bacterial growth by the innate immune response was needed to recapitulate the threshold effect associated with the effective bacterial inoculum. This phenomenon could be well captured using our model (Eq. 3), with estimates of *δ_M_*, the maximal decay induced by the immune response, of 6.8 log_10_ CFUeq/g/h and *B*_50_, the bacterial load needed to reach *δ_M_*/2, of 7.4 log_10_ CFUeq/g. This corresponds to a threshold for bacterial control equal to 6.94 log_10_ CFUeq/g. Thus, if the administered bacterial inoculum, *B*_0_, was below this value, bacteria were rapidly eliminated by the immune system with a rate ~ *δ_M_/B*_50_ = 0.3 h^−1^, corresponding to a half-life of 2.3 hours. However, when the bacterial inoculum was above this value, the model predicts a saturation of the immune response, enabling bacterial growth with a bacterium doubling time estimated at 3.5 hours.

We performed *in vitro* experiments from which we estimated two key parameters of phage dynamics that could not be identified *in vivo,* namely the median eclipse phase and the burst size (see methods and Supplementary Figure S2). Assuming that these parameters had the same value *in vivo,* we could estimate the infection rate of bacteria by phages, noted *β,* to 6.9 log_10_ g/CFU/h, and the elimination rate of productively infected bacteria, *δ_V_,* to 0.09 h^−1^ (eg, a half-life of 7.7 h).

Finally, we also used the model to assess the impact of bacterial kinetics on survival. Assuming a log-linear effect of bacterial load on the instantaneous risk of death, *h(t),* the bacterial load was found to be highly associated with death (*γ*=1.44, p<10^−3^, Wald test). This corresponds to a hazard ratio of 4.2; in other words, an increase of 2 log of the bacterial load raised the instantaneous risk of death by ~18. The joint model developed could simultaneously recapitulate bacterial kinetics, phage counts and survival rate (Supplementary Figures S5-S6), despite the heterogeneity of the individual profiles experimentally observed (Figure 5).

**Figure 5.**
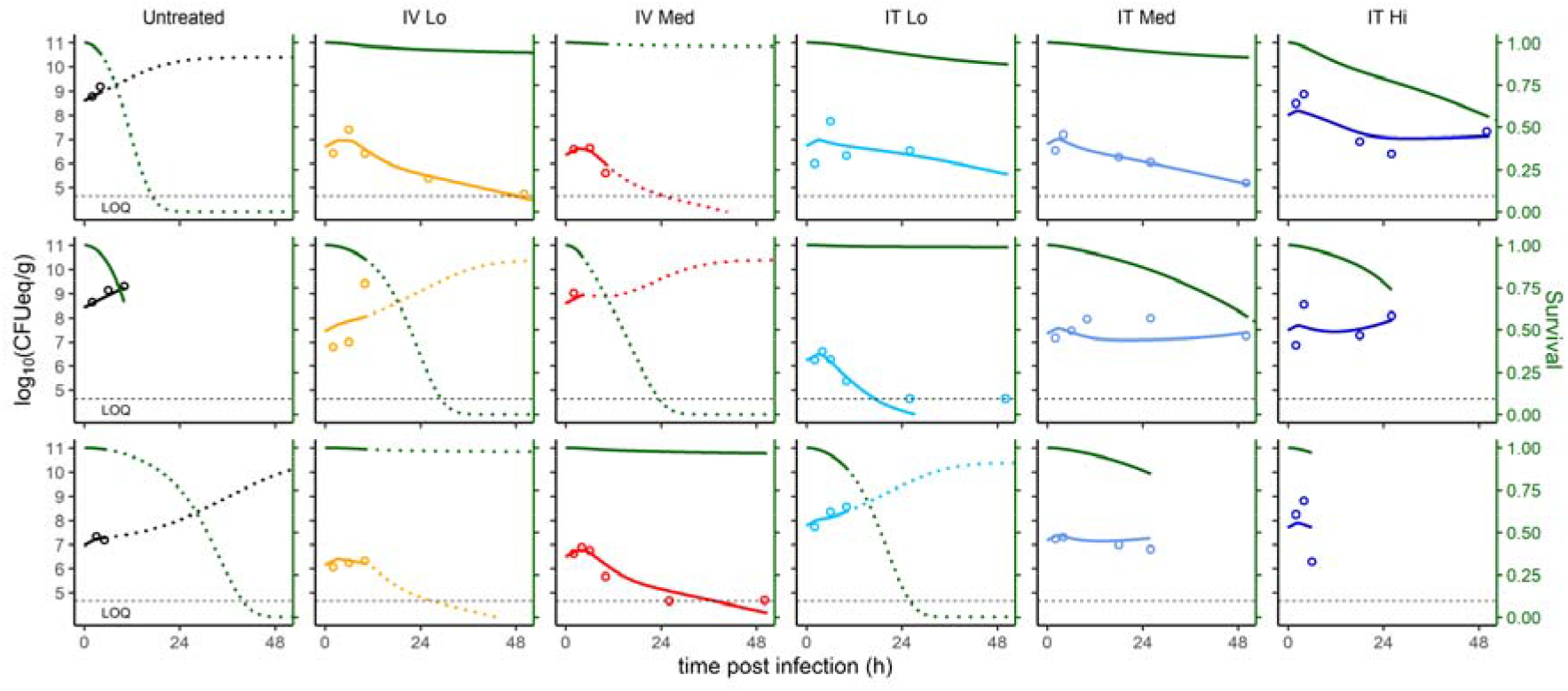
The mathematical model recapitulates the individual variations of the pulmonary bacterial loads and estimates the survival of infected animals. Data and model predictions for three mice (lines) randomly chosen from each experimental groups (columns) are shown. The circles correspond to the pulmonary bacterial loads, the color lines correspond to the individual predictions from the model, and the green line represents the predicted survival probability. Lines are solid between t=0 and the last observation time point (death or pre-specified sacrifice) and are dotted after the pre-specified sacrifice. Interrupted lines indicate the death of animals before the pre-specified time point. The horizontal dashed line represents the limit of quantification (LOQ=4.7 log_10_ CFUeq/g).

### Effects of bacterial inoculum, dose, route of administration and phage characteristics on the bacterial kinetics

The model was then used to identify the key factors governing bacterial kinetics. Unsurprisingly, the bacterial inoculum, *B_0_,* was the most important parameter. In animals with B_0_<6 log_10_ CFUeq/g, bacterial loads rapidly decreased below the limit of quantification within 48h, regardless of treatment. In animals with large bacterial inoculum B_0_>8 log_10_ CFUeq/g, phage therapy was not sufficient to control the continuous growth of bacteria, regardless of the route and administered dose (Figure 5, Figure 6). The benefit of phage therapy was visible mostly for animals having an intermediate effective inoculum size, e.g. 6<B_0_<8 log_10_ CFUeq/g, in which phages were critical to prevent bacteria growth and to maintain the bacterial load under control by the immune system. In those animals, the decline of bacterial load was biphasic, with a rapid decline in the first four hours of treatment, followed by a slower decline afterwards. This slower second phase was attributed in our model to the unrestricted replication of a sub-population of bacteria that were not eliminated by phages, representing both phage-resistant and phage-inaccessible (spatially out of reach) bacteria. The initial proportion of this phage-refractory population was estimated to 10% (Table 1), but could be variable across individuals (from 0.03% to 76%). The dynamics of refractory bacteria was then individual dependent, leading either to a slower but continuous bacterial decline or to a rebound of fully refractory bacteria within 24-48 hours (see individual examples in Figure 5). Importantly, the need to incorporate a compartment of refractory bacteria in the model was strongly supported by data and led to a significant improvement of the data fitting (p<10^−3^, Likelihood Ratio Test, not shown).

**Figure 6.**
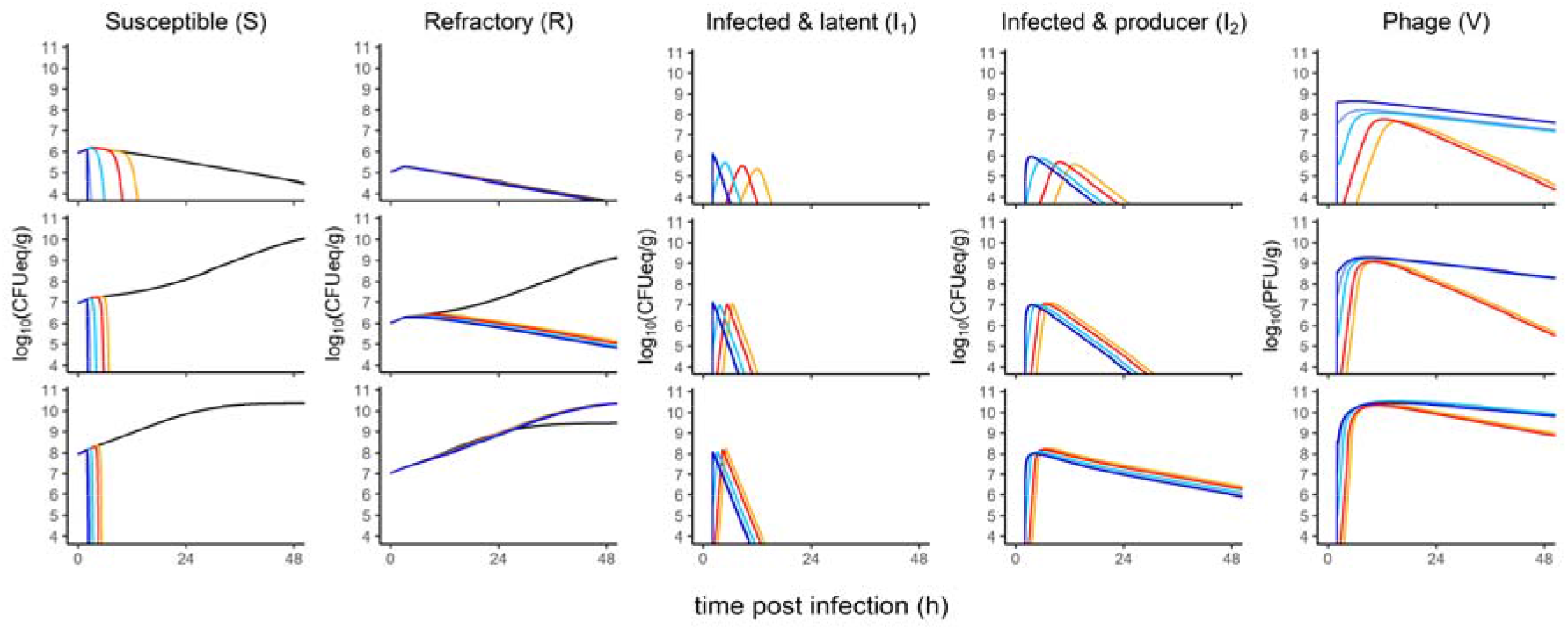
Simulations of different bacterial inoculums inform on the impact of refractory bacteria. Median profile assuming initial bacterial inoculums of 6, 7 and 8 log_10_ CFUeq/g (top, middle and bottom lines, respectively). Predictions were calculated for each bacterial population (S, R, I1 and I2) and phages (V). The colors are indicative of both the route of administration (IT or IV) and the dose of phage received (Lo: 5.6 log_10_ PFU; Med: 7.6 log_10_ PFU; Hi: 8.6 log_10_ PFU): untreated (black), IV-Lo (orange), IV-Med (red), IT-Lo (cyan), IT-med (blue), IT-high (dark blue).

The raw data did not allow to observe a role of route of administration in treatment outcome, except in the case of a low bacterial inoculum (B_0_=6 log_10_ CFUeq/g) in which the rapid clearance of bacteria limited the encounter rate with phage. Our simulations indicated that IT and IV otherwise reached similarly high phages concentrations in lungs (Figure 6). IT administration was however associated with a substantially faster time to peak viral load, and this difference was particularly obvious with a low or an intermediate bacterial inoculum. For instance, with an intermediate bacterial inoculum (*B*_0_=7 log_10_ CFUeq/g), the time to peak viral load varied between 1.6 h (IT high dose) and 6.0 h (IV low dose). As a consequence of this delay in phage maximal replication, IV route had a less efficiency to stop bacterial growth, with a time to peak bacterial load that could be as large as 4.0 h (IV low dose), to less than 1.1 h with IT route, and as low as 0.3 h in case of IT high dose.

Finally, we also used the model to explore the effect of intrinsic phage characteristics, namely burst size and lysis rate, on the bacterial kinetics. A 10-fold reduction in burst size *(i.e.* 57 PFU/CFU) predicted a larger impact of the route of administration, with a time to bacterial peak of 10 h post-IV compared to less than 3 h post-IT administration, even when using a low dose (Figure S7). However, a higher lysis rate value counter-balanced the effects of a low burst size value (Figure S8). These showed the sensitivity of the model to phage characteristics.

## Discussion

Despite many encouraging data from both ancient literature and recent compassionate treatments, phage therapy remains underexploited, notably because of ambiguous results from clinical trials (Luong et al., 2020). Indeed, the translation from *in vitro* to clinical conditions remains unmet, in particular because using phages as a drug requires a specific method to elaborate a rational scheme for safe and optimal administration. While some studies performed *in silico* or in laboratory conditions (Cairns et al., 2009; Kasman et al., 2002; Leung and Weitz, 2017; Payne and Jansen, 2001, 2003) led to propose several mathematical models of phage-bacteria interactions, very few studies including the host immune response were conducted *in vivo* (Dabrowska and Abedon, 2019; Roach et al., 2017). Here, we developed a pharmacometric model, exploiting *in vivo* data from a murine pneumonia treated by a single virulent phage administered at three doses and via two administration routes, which robustness could support further the (re)deployment of phage therapy.

Our experimental approach was inspired from the work performed during the development of new drugs, where intensive pharmacokinetic and pharmocodynamic data are required to optimize the dosing regimens of a treatment (Marshall et al., 2019). Here, we collected an unprecedented biological set of data using an *in vivo* imaging tool to longitudinally follow the same animals over time and from which we developed a semi-mechanistic model to characterize the dynamics of a phage treatment.

The biodistribution of phages in the lungs of uninfected mice showed that phage loads were much higher and their exposition lasted longer following local (IT) compared to systemic (IV) administration. Consequently, IT administration led to a rapid decline of bacterial loads within the next two hours post-administration. Nevertheless, 12 hours post local and systemic phage administrations, all dosing regimens led to comparable levels of phage exposure, due to the large burst size of phage 536_P1. Despite being in agreement with previous literature (Morello et al., 2011; Oechslin et al., 2017), these biodistribution data cannot be yet generalized to all phages given the very large diversity of these viruses. Also, we did not study repeated IV administered doses that could allow for longer and higher phage exposition in lungs.

The heterogeneity of the initial bacterial loads observed two hours post-infection explains the lack of statistical differences in Kaplan-Meir survival estimations when pooling mice to compare routes of administration and doses. The source of this heterogeneity was the consequence of choices made to maximize the probability of interactions between phages and bacteria, compared to our previous work using the same couple of phage-bacteria (Dufour et al., 2015). Here, we increased the inoculum and we used the direct intratracheal route (instead of the intranasal route) to reach the lung, in order to reduce spatial dispersion along the upper respiratory tract. The large range of situations observed was instrumental to enrich the mathematical model with multiple events (N=236 analyzed mice) to improve the assessments of phage-bacteria parameters, doses and routes of administration. Moreover, a direct consequence of the above conditions was the precise quantification of the threshold value (6.94 log_10_ CFUeq/g) above which the immune response was overwhelmed, which was very close to the value (6.80 log_10_ CFUeq/g) deduced from our model.

The modeling of phage-bacteria interactions in the host required the definition of a refractory bacterial population. This compartment could not be phenotypically characterized but encompasses bacteria that are escaping phage predation because they are phage-resistant and/or they remain spatially inaccessible to phages (Leung and Weitz, 2017). The inclusion of this compartment is supported by both the emergence of phage resistance observed *in vitro* after 10 h of incubation with phage 536_P1 (Supplementary Figure S2), and previous experimental and theoretical studies showing that the spatial distribution of bacteria in organs can prevent bacteria to be infected by phages (Lourenco et al., 2020; Sousa and Rocha, 2019). This was also in agreement with the modelling of phage therapy in immunocompromised hosts (Roach et al., 2017). Despite the large variation of the bacterial loads that directly affects the initial density of this refractory population, our model could appropriately recapitulate the kinetics of phage-bacteria interactions for each individual mouse, including the survival probability, testifying of its robustness. In addition, simulations performed with larger or lower burst sizes and lysis rates, showed that the model predicts outputs that are in agreement with results from animal studies in which different phages were used to treat bacterial infections (Alemayehu et al., 2012; Carmody et al., 2010; Henry et al., 2013; Morello et al., 2011).

To ensure the model recapitulates kinetics of phage-bacteria interactions during therapy, it was necessary to use as much as possible parameters deduced from animal experiments instead of *in vitro* conditions. For instance, growth rate of bacteria, infectivity rate and lysis rate, were all deduced from experimental data. Nevertheless, other parameters should be more deeply investigated to improve the fidelity of the model. For instance, growth of refractory population was approximated as similar to the susceptible bacteria, neglecting the possibility that the fitness of mutants might be affected in the host. Indeed, such mutants could be, either or both, less virulent and strongly affected in their growth, allowing immune cells to eliminate them more effectively (Oechslin et al., 2017; Pouillot et al., 2012; Smith et al., 1987). The involvement of the immune response in the success of respiratory phage therapy was previously demonstrated and modelled, but remains mechanistically poorly explored (Roach et al., 2017). In this work we chose to account for the immune response in a synthetic way which encompasses mechanical factors, such as mucins and surfactant proteins, as well as various cellular actors of the innate immune system (Bergeron et al., 1998; Marriott and Dockrell, 2007).

Predictions from the mathematical model showed the importance of viral load, burst size and lysis rate to fully capture bacterial kinetics and animal outcome. Therefore, in a clinical context, the administration route, and to a lesser degree the dose, will play a critical role in the rate at which bacterial growth will be controlled, supporting the determination of key parameters of phage-bacteria interactions as a prerequisite for optimizing phage therapy. However, the model predicted that substantial variations of the intrinsic characteristics of phages (burst size and lysis rate) were not jeopardizing treatment output, which fits with the current semi-empirical choice of phages for compassionate treatments. Indeed, currently phages are chosen exclusively for their capacity to infect bacteria on *in vitro* conditions where phage-bacteria interactions are artificially optimized. The model is therefore robust, which argues in favor of its relevance for many other phages.

To conclude, the model developed during this work could be used to predict the efficacy of virtually any phage for which a minimum set of *in vitro* and *in vivo* data must be obtained, which will considerable lower the number of experiments needed to validate such a phage during pre-clinical development. Beyond phage therapy, the model could also be implemented to test combined anti-infectious therapies such as the association of phages with antibiotics (Lin et al., 2020).

## Methods

### Determination of *in vitro* infection parameters of phage 536_P1

Parameters of the interactions between strain 536 and phage 536_P1 have been determined *in vitro,* notably the rate of transition *e* from infected non-productive bacteria to productively infected bacteria (eg, log(2)/e is the median eclipse phase duration), and the burst size *b, i.e.* the number of virions released by a lysed bacteria. The median eclipse phase time and the burst size were estimated from non-linear mixed effect models performed with the SAEM algorithm implemented in Monolix software (version 2018R2) (Kuhn and Lavielle, 2005) using data from one-step-growth experiments (see Supplementary material).

### Presentation of the murine model of pulmonary phage therapy

The *E. coli* strain 536 is a virulent clinical strain belonging to the B2 phylogenetic group (Brzuszkiewicz et al., 2006). The *E. coli* strain 536-lux was obtained by the transformation of wild-type 536 strain with plasmid pCM17, which carries the *luxCDABE* operon from *Photorhabdus luminescens* under *OmpC* promoter (Morin and Kaper, 2009). This plasmid is stable over 7 days without adding antibiotic in the medium (Dufour et al., 2015).

Phage 536_P1 (Dufour et al., 2015) is a virulent *Myoviridae* (149.4 Kb) that was amplified on strain 536 and purified according to molecular biology protocols including an endotoxin removal step (Endotrap Blue, Hyglos, Germany). The stock solution of 8.2 log_10_ PFU/mL has an endotoxin concentration of 2.15 EU/mL.

Pneumonia was induced by intratracheal (IT) administration of strain 536-lux in eight-week-old BALB/cJRj anesthetized male mice (mixture of Ketamine and Xylazine administered intraperitoneally, IP). Exponentially growing bacteria in Luria Bertoni Broth (Lennox) at 37°C were centrifuged and washed twice and finally resuspended in PBS to obtain inoculums ranging from 7.6 to 8.2 log_10_ CFU in 20μL.

Phage treatment was applied under general anesthesia (isoflurane, inspired fraction=2%) using IT or intravenous (IV, retro-orbital injection) administration two hours after infection. A volume of 20μL of phage 536_P1 was used, containing either a low (Lo; 6.6 log_10_ PFU), medium (Me; 7.6 log_10_ PFU) or high (Hi; 8.6 log_10_ PFU) amount of phages.

At 2, 4, 8, 24 and 48h after phage administration, mice were sacrificed, and blood (in heparin coated tubes) and lungs were recovered and put on ice. Lungs were weighted, homogenized in PBS with protease inhibitor (EDTA free micro Tabs, SigmaAldrich). Lungs and blood were then centrifuged (5000g, 15 min, 4°C). The supernatant was collected, serially diluted and spotted (in triplicate) on agar plates covered with strain 536 to titer phage 536_P1.

### Bioluminescence and longitudinal follow up of bacterial load in lungs

In all our experiments photon emission of the luminescent bacteria was recorded at regular intervals under general anesthesia (isoflurane, inspired fraction=2%) using IVIS Spectrum and analyzed with Living Image software (PerkinElmer, Richmond, CA) as previously described (Debarbieux et al., 2010). Photons emitted within the chest area of each mouse were collected as photon/s/cm^2^/steradian.

To fully exploit the *in vivo* imaging tool we established the correlation between bioluminescence and CFU count in lungs using data from 17 untreated animals that were euthanized at different time points and for which lung bacteria was quantified by plating (Figure S1). The relationship obtained allowed us to derive from the luminescence signal an equivalent counts of CFU per gram of lung (noted CFUeq/g) that was used to obtain in all infected animals (untreated or treated by phages) the bacterial densities.

### Phage distribution

A dose of 8 log_10_ PFU of phage 536_P1 was administered in uninfected mice (N=53) using IT (N=16) or IV (N=37) route. At 2, 4, 8, 24 and 48h after phage administration, mice were sacrificed, and blood (in heparin coated tubes) and lungs were recovered and put on ice. Lungs were weighted, homogenized in PBS with protease inhibitor (EDTA free micro Tabs, SigmaAldrich). Lungs and blood were then centrifuged (5000g, 15 min, 4°C). The supernatant was collected, serially diluted and spotted (in triplicate) on agar plates covered with strain 536 to titer phage 536_P1.

We used the following standard pharmacokinetic model to estimate phage biodistribution:

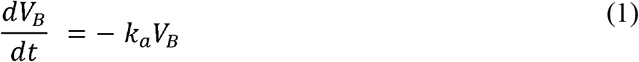

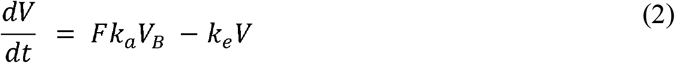

Where *V* and *V_B_* denote phage quantity in lung and blood compartments, respectively, *k_e_* is the elimination rate from lungs, *k_a_* is the transfer rate from blood to lungs, *F* is the bioavailability in lungs after IV administration. The initial condition of these equations are *V_B_*(0) = *dose* and *V*(0) = 0 in case of IV administration and *V_B_*(0) = 0 and *V*(0) = *dose* in case of IT administration. *V* is divided by the phage distribution weight (*W*) to obtain the phage concentration in lung compartment. The parameter *W* was deduced from the extrapolation of the median phage concentration at baseline (8 log_10_ PFU/g) following the IT administration of a single dose of 8 log_10_ PFU, and F was obtained by performing the ratio of IV/IT area under the curves in the lung. Estimations of the other parameters were performed by naïve pooling of data (i.e. considering all the measures for each group (IV and IT) as if they came from a single individual and performing a nonlinear regression).

### Bacterial kinetics in untreated mice

We characterize the natural bacterial kinetics (i.e., in absence of treatment) in animals infected with doses of 7.6 log_10_ CFU (34 mice) or 8.2 log_10_ CFU (12 mice) and the bacterial kinetics obtained by bioluminescence (see above) using the following model (one mouse was not analyzed due to death during bacterial inoculation, i.e. N=45):

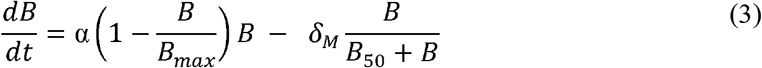

Where *α* is the exponential growth rate of bacteria (eg, log(2)/*α* is the doubling time) and *B_max_* is the maximal carrying capacity. The initial condition of *B* was estimated by the parameter *B*_0_ (*B*(0) = 10^*B*_o_^). Consistent with previous work (Dufour et al., 2019), we assumed that in the first 3 hours of infection the immune system had no activity (eg, bacteria could grow exponentially during this period). After this period the elimination of bacteria by the host was modeled by a non-linear saturable term, given by 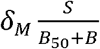. The model parameters were estimated in non-linear mixed effect models performed with the SAEM algorithm implemented in Monolix software (version 2018R2) (Kuhn and Lavielle, 2005).

### Phage-bacteria interaction in treated mice

We characterized phage/bacterial interactions using data obtained in 142 mice infected by 7.6 log_10_ CFU and treated subsequently (four mice were not analyzed due to death during bacterial inoculation, i.e. N=138), using the protocol described above, with a single dose (low, medium or high) of phages administered using IT or IV route (detail of the groups in Figure 1). Bacterial kinetics was derived from bioluminescence (see above) and lungs were collected at 2, 4, 8, 24 or 48h after phage administration to count PFU. We also considered data in untreated mice (see previous section) for the analysis.

The model describing interactions between phage 536_P1 and strain 536 relied on a system of ordinary differential equations (ODE) (Figure 1, Equations 4 to 10, presented below). We combined standard model of viral (Perelson and Guedj, 2015) and bacterial kinetics (Friberg and Guedj, 2020) to tease out the main parameters governing phage-bacteria interaction in mice. In lungs, phages can infect susceptible bacteria (S) following a mass-action term, with rate *β.* We also considered a population of refractory bacteria (of initial proportion *f*), noted R, that includes both phage-resistant and phage-inaccessible bacteria.

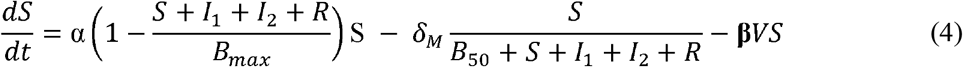

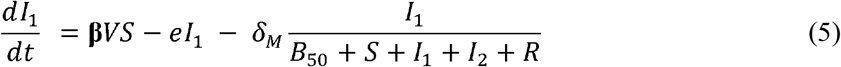

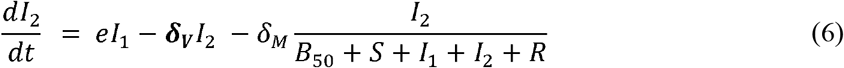

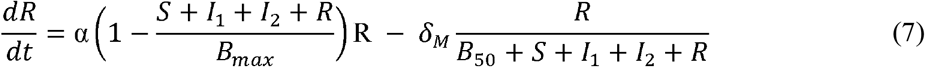

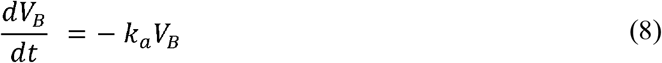

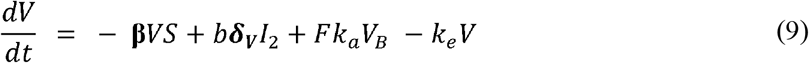

We considered the following initial condition for this system: *S*(0) = 10^*B*_0_^(1 – *f*), *l*_1_(0) = 0, *I*_2_(0) = 0, *R*(0) = 10^*B*_0_^ × ***f***, *V_B_*(0) = 0, *V*(0) = θ. The dose of phages is added to *V* or *V_B_* at the phage administration time by IT or IV route, respectively.

The model parameters in bold were estimated in non-linear mixed effect models performed with the SAEM algorithm implemented in Monolix software using data from all infected animals (version 2018R2) (Kuhn and Lavielle, 2005). All other parameters were fixed to the values determined above, either *in vitro* (*e,b*) in uninfected treated mice (*k_a_, F, k_e_*) or in infected untreated mice (*α, B_max_, δ_M_, β*_50_).

### Impact of bacterial kinetics on survival

Finally, we aimed to take into account the potential bias due to the fact that more severely infected animals died earlier, making the population study less and less representative as time goes by. In this approach, called joint models (Desmee et al., 2015), the probability to survive up to time *t, S*(*t*), is modeled as: 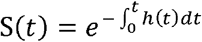 where

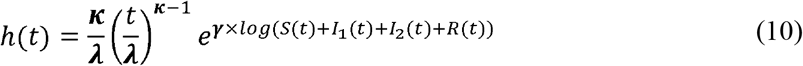

h(t) is the instantaneous hazard function, *κ* and *λ* are scale parameters characterizing the risk of death in absence of infection, and *γ* quantifies the impact of bacterial load on mortality. Thus, in this model *γ* = 0 is indicative of an absence of association between bacterial load and mortality, while *γ >* 0 indicates that increased bacterial load is associated with an increased risk of death.

### Model evaluation

Individual fits were investigated to evaluate the different models. For the final model, we also evaluated the model predictions i) for the bacterial kinetics by using prediction corrected-visual predictive checks (Bergstrand et al., 2011), accounting for the risk of death and ii) visual predictive checks of the survival. Simulations for validation methods were made using mlxR package.

### Model based simulations for the impact of treatment procedures and phage parameter values

Model based predictions were performed to investigate various scenarios as follows. For each simulation, the median profile of different model compartments (*S,R,I*_1_, *I*_2_ and *V*) was investigated using the parameters determined in Table 1 and different experimental conditions regarding the route of administration (IV or IT), the dose of phage administered (Lo: 5.6, Med: 7.6 or Hi: 8.6 log_10_ PFU) and initial bacterial load (6, 7 or 8 log_10_ CFUeq/g). We also evaluated the effects of different burst size (10 fold lower, 10 fold larger than the estimated value *in vitro*), using a single initial bacterial load of 7 log_10_ CFUeq/g. Furthermore, when setting a burst size 10 times lower than the estimated value (and a fixed initial bacterial load of 7 log_10_ CFUeq/g), we investigated the effect of three different lysis rates values: the value estimated *in vivo* (0.09 h^−1^); a value three times higher (0.27 h^−1^); a value estimated *in vitro* (1.39 h^−1^).

### Ethical statement

A total of 241 eight-week-old BALB/cJRj male mice (Janvier, France) were housed in animal facility in accordance with French and European regulations on the care and protection of laboratory animals. Food and drink were provided ad libitum. Protocol was approved by the veterinary staff of the Institut Pasteur animal facility (approval number 20.173) and the National Ethics Committee regulating animal experimentation (APAFIS#26874-2020081309052574 v1).

## Supporting information

Supplementary content

## Authors contribution

JDR, JG, LD and RD, conceived the study and secured funding. BG, FH, RD and RK performed experiments. JDR, JG, JS, LD and RD analyzed the data. JG, JS and TTN performed modeling. ND provided feedback. JS and RD wrote the first draft. JG and LD reviewed and edited final draft.

## Declaration of interests

Authors declare no competing interest.

## Notes

### Competing Interest Statement

The authors have declared no competing interest.

## References

Alemayehu, D., Casey, P.G., McAuliffe, O., Guinane, C.M., Martin, J.G., Shanahan, F., Coffey, A., Ross, R.P., and Hill, C. (2012). Bacteriophages phiMR299-2 and phiNH-4 can eliminate Pseudomonas aeruginosa in the murine lung and on cystic fibrosis lung airway cells. mBio 3, e00029–00012.

Aslam, S., Lampley, E., Wooten, D., Karris, M., Benson, C., Strathdee, S., and Schooley, R.T. (2020). Lessons Learned From the First 10 Consecutive Cases of Intravenous Bacteriophage Therapy to Treat Multidrug-Resistant Bacterial Infections at a Single Center in the United States. Open forum infectious diseases 7, ofaa389.

Bergeron, Y., Ouellet, N., Deslauriers, A.M., Simard, M., Olivier, M., and Bergeron, M.G. (1998). Cytokine kinetics and other host factors in response to pneumococcal pulmonary infection in mice. Infect Immun 66, 912–922.

Bergstrand, M., Hooker, A.C., Wallin, J.E., and Karlsson, M.O. (2011). Prediction-corrected visual predictive checks for diagnosing nonlinear mixed-effects models. AAPS J 13, 143–151.

Brzuszkiewicz, E., Bruggemann, H., Liesegang, H., Emmerth, M., Olschlager, T., Nagy, G., Albermann, K., Wagner, C., Buchrieser, C., Emody, L., et al. (2006). How to become a uropathogen: comparative genomic analysis of extraintestinal pathogenic Escherichia coli strains. Proc Natl Acad Sci U S A 103, 12879–12884.

Cairns, B.J., Timms, A.R., Jansen, V.A., Connerton, I.F., and Payne, R.J. (2009). Quantitative models of in vitro bacteriophage-host dynamics and their application to phage therapy. PLoS Pathog 5, e1000253.

Carmody, L.A., Gill, J.J., Summer, E.J., Sajjan, U.S., Gonzalez, C.F., Young, R.F., and LiPuma, J.J. (2010). Efficacy of bacteriophage therapy in a model of Burkholderia cenocepacia pulmonary infection. J Infect Dis 201, 264–271.

Corbellino, M., Kieffer, N., Kutateladze, M., Balarjishvili, N., Leshkasheli, L., Askilashvili, L., Tsertsvadze, G., Rimoldi, S.G., Nizharadze, D., Hoyle, N., et al. (2020). Eradication of a Multidrug-Resistant, Carbapenemase-Producing Klebsiella pneumoniae Isolate Following Oral and Intra-rectal Therapy With a Custom Made, Lytic Bacteriophage Preparation. Clin Infect Dis 70, 1998–2001.

Dabrowska, K., and Abedon, S.T. (2019). Pharmacologically Aware Phage Therapy: Pharmacodynamic and Pharmacokinetic Obstacles to Phage Antibacterial Action in Animal and Human Bodies. Microbiol Mol Biol Rev 83.

Debarbieux, L., Leduc, D., Maura, D., Morello, E., Criscuolo, A., Grossi, O., Balloy, V., and Touqui, L. (2010). Bacteriophages can treat and prevent Pseudomonas aeruginosa lung infections. J Infect Dis 201, 1096–1104.

Desmee, S., Mentre, F., Veyrat-Follet, C., and Guedj, J. (2015). Nonlinear Mixed-effect Models for Prostate-specific Antigen Kinetics and Link with Survival in the Context of Metastatic Prostate Cancer: A Comparison by Simulation of Two-stage and Joint Approaches. AAPS J 17, 691–699.

Dion, M.B., Oechslin, F., and Moineau, S. (2020). Phage diversity, genomics and phylogeny. Nat Rev Microbiol 18, 125–138.

Dufour, N., Debarbieux, L., Fromentin, M., and Ricard, J.D. (2015). Treatment of Highly Virulent Extraintestinal Pathogenic Escherichia coli Pneumonia With Bacteriophages. Crit Care Med 43, e190–198.

Dufour, N., Delattre, R., Chevallereau, A., Ricard, J.D., and Debarbieux, L. (2019). Phage Therapy of Pneumonia Is Not Associated with an Overstimulation of the Inflammatory Response Compared to Antibiotic Treatment in Mice. Antimicrob Agents Chemother 63.

Friberg, L.E., and Guedj, J. (2020). Acute bacterial or viral infection-What’s the difference? A perspective from PKPD modellers. Clin Microbiol Infect 26, 1133–1136.

Gordillo Altamirano, F.L., and Barr, J.J. (2019). Phage Therapy in the Postantibiotic Era. Clin Microbiol Rev 32.

Henry, M., Lavigne, R., and Debarbieux, L. (2013). Predicting in vivo efficacy of therapeutic bacteriophages used to treat pulmonary infections. Antimicrob Agents Chemother 57, 5961–5968.

Jault, P., Leclerc, T., Jennes, S., Pirnay, J.P., Que, Y.A., Resch, G., Rousseau, A.F., Ravat, F., Carsin, H., Le Floch, R., et al. (2019). Efficacy and tolerability of a cocktail of bacteriophages to treat burn wounds infected by Pseudomonas aeruginosa (PhagoBurn): a randomised, controlled, double-blind phase 1/2 trial. Lancet Infect Dis 19, 35–45.

Jennes, S., Merabishvili, M., Soentjens, P., Pang, K.W., Rose, T., Keersebilck, E., Soete, O., Francois, P.M., Teodorescu, S., Verween, G., et al. (2017). Use of bacteriophages in the treatment of colistin-only-sensitive Pseudomonas aeruginosa septicaemia in a patient with acute kidney injury-a case report. Crit Care 21, 129.

Jiang, W., Yan, Y., Ji, W., Wang, Y., and Chen, Z. (2014). Clinical significance of different bacterial load of Mycoplasma pneumoniae in patients with Mycoplasma pneumoniae pneumonia. Braz J Infect Dis 18, 124–128.

Kasman, L.M., Kasman, A., Westwater, C., Dolan, J., Schmidt, M.G., and Norris, J.S. (2002). Overcoming the phage replication threshold: a mathematical model with implications for phage therapy. J Virol 76, 5557–5564.

Kortright, K.E., Chan, B.K., Koff, J.L., and Turner, P.E. (2019). Phage Therapy: A Renewed Approach to Combat Antibiotic-Resistant Bacteria. Cell Host Microbe 25, 219–232.

Kuhn, E., and Lavielle, M. (2005). Maximum likelihood estimation in nonlinear mixed effects models. Computational Statistics & Data Analysis 49, 1020–1038.

Leitner, L., Ujmajuridze, A., Chanishvili, N., Goderdzishvili, M., Chkonia, I., Rigvava, S., Chkhotua, A., Changashvili, G., McCallin, S., Schneider, M.P., et al. (2020). Intravesical bacteriophages for treating urinary tract infections in patients undergoing transurethral resection of the prostate: a randomised, placebo-controlled, double-blind clinical trial. Lancet Infect Dis.

Leung, C.Y.J., and Weitz, J.S. (2017). Modeling the synergistic elimination of bacteria by phage and the innate immune system. J Theor Biol 429, 241–252.

Lin, Y., Quan, D., Yoon Kyung Chang, R., Chow, M.Y.T., Wang, Y., Li, M., Morales, S., Britton, W.J., Kutter, E., Li, J., et al. (2020). Synergistic activity of phage PEV20-ciprofloxacin combination powder formulation-A proof-of-principle study in a P. aeruginosa lung infection model. European journal of pharmaceutics and biopharmaceutics: official journal of Arbeitsgemeinschaft fur Pharmazeutische Verfahrenstechnik eV.

Lourenco, M., Chaffringeon, L., Lamy-Besnier, Q., Pedron, T., Campagne, P., Eberl, C., Berard, M., Stecher, B., Debarbieux, L., and De Sordi, L. (2020). The Spatial Heterogeneity of the Gut Limits Predation and Fosters Coexistence of Bacteria and Bacteriophages. Cell Host Microbe 28, 390–401 e395.

Luong, T., Salabarria, A.C., and Roach, D.R. (2020). Phage Therapy in the Resistance Era: Where Do We Stand and Where Are We Going? Clin Ther 42, 1659–1680.

Marriott, H.M., and Dockrell, D.H. (2007). The role of the macrophage in lung disease mediated by bacteria. Exp Lung Res 33, 493–505.

Marshall, S., Madabushi, R., Manolis, E., Krudys, K., Staab, A., Dykstra, K., and Visser, S.A.G. (2019). Model-Informed Drug Discovery and Development: Current Industry Good Practice and Regulatory Expectations and Future Perspectives. CPT: pharmacometrics & systems pharmacology 8, 87–96.

Melo, L.D.R., Oliveira, H., Pires, D.P., Dabrowska, K., and Azeredo, J. (2020). Phage therapy efficacy: a review of the last 10 years of preclinical studies. Crit Rev Microbiol 46, 78–99.

Morello, E., Saussereau, E., Maura, D., Huerre, M., Touqui, L., and Debarbieux, L. (2011). Pulmonary bacteriophage therapy on Pseudomonas aeruginosa cystic fibrosis strains: first steps towards treatment and prevention. PLoS One 6, e16963.

Morin, C.E., and Kaper, J.B. (2009). Use of stabilized luciferase-expressing plasmids to examine in vivo-induced promoters in the Vibrio cholerae vaccine strain CVD 103-HgR. FEMS Immunol Med Microbiol 57, 69–79.

Oechslin, F., Piccardi, P., Mancini, S., Gabard, J., Moreillon, P., Entenza, J.M., Resch, G., and Que, Y.A. (2017). Synergistic Interaction Between Phage Therapy and Antibiotics Clears Pseudomonas Aeruginosa Infection in Endocarditis and Reduces Virulence. J Infect Dis 215, 703–712.

Payne, R.J., and Jansen, V.A. (2001). Understanding bacteriophage therapy as a density-dependent kinetic process. J Theor Biol 208, 37–48.

Payne, R.J., and Jansen, V.A. (2003). Pharmacokinetic principles of bacteriophage therapy. Clin Pharmacokinet 42, 315–325.

Perelson, A.S., and Guedj, J. (2015). Modelling hepatitis C therapy--predicting effects of treatment. Nat Rev Gastroenterol Hepatol 12, 437–445.

Pouillot, F., Chomton, M., Blois, H., Courroux, C., Noelig, J., Bidet, P., Bingen, E., and Bonacorsi, S. (2012). Efficacy of bacteriophage therapy in experimental sepsis and meningitis caused by a clone O25b:H4-ST131 Escherichia coli strain producing CTX-M-15. Antimicrob Agents Chemother 56, 3568–3575.

Roach, D.R., Leung, C.Y., Henry, M., Morello, E., Singh, D., Di Santo, J.P., Weitz, J.S., and Debarbieux, L. (2017). Synergy between the Host Immune System and Bacteriophage Is Essential for Successful Phage Therapy against an Acute Respiratory Pathogen. Cell Host Microbe 22, 38–47 e34.

Sarker, S.A., Sultana, S., Reuteler, G., Moine, D., Descombes, P., Charton, F., Bourdin, G., McCallin, S., Ngom-Bru, C., Neville, T., et al. (2016). Oral Phage Therapy of Acute Bacterial Diarrhea With Two Coliphage Preparations: A Randomized Trial in Children From Bangladesh. EBioMedicine 4, 124–137.

Schooley, R.T., Biswas, B., Gill, J.J., Hernandez-Morales, A., Lancaster, J., Lessor, L., Barr, J.J., Reed, S.L., Rohwer, F., Benler, S., et al. (2017). Development and Use of Personalized Bacteriophage-Based Therapeutic Cocktails To Treat a Patient with a Disseminated Resistant Acinetobacter baumannii Infection. Antimicrob Agents Chemother 61.

Smith, A.M., McCullers, J.A., and Adler, F.R. (2011). Mathematical model of a three-stage innate immune response to a pneumococcal lung infection. J Theor Biol 276, 106–116.

Smith, H.W., Huggins, M.B., and Shaw, K.M. (1987). The control of experimental Escherichia coli diarrhoea in calves by means of bacteriophages. J Gen Microbiol 133, 1111–1126.

Sousa, J.A.M., and Rocha, E.P.C. (2019). Environmental structure drives resistance to phages and antibiotics during phage therapy and to invading lysogens during colonisation. Sci Rep 9,3149.

Weld, R.J., Butts, C., and Heinemann, J.A. (2004). Models of phage growth and their applicability to phage therapy. J Theor Biol 227, 1–11.

