## Supplementary content for "Combination of *in vivo* phage therapy data with *in silico* model highlights key parameters for treatment efficacy"

by Delattre et al

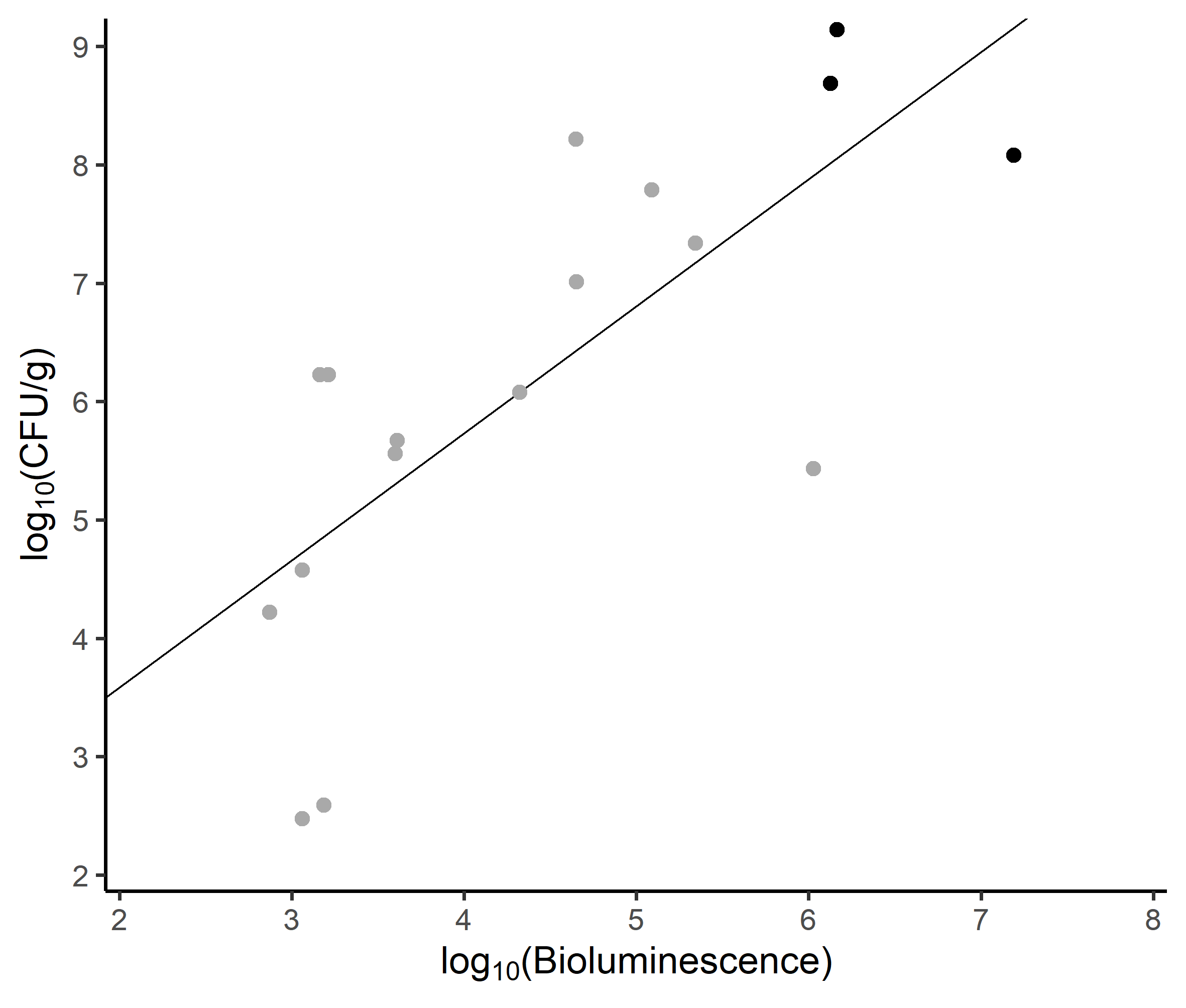

**Supplementary figure S1.** Correlation between bioluminescence and CFU from lungs of infected and untreated mice. The regression line equation is $\log_{10} (CFU/g)=1.44+1.07\log_{10} (Bioluminescence)$. From 17 mice, 14 were infected with an inoculum of 7.6 log_10_ CFU (grey dots) and 3 with an inoculum of 8.2 log_10_ CFU (black dots).

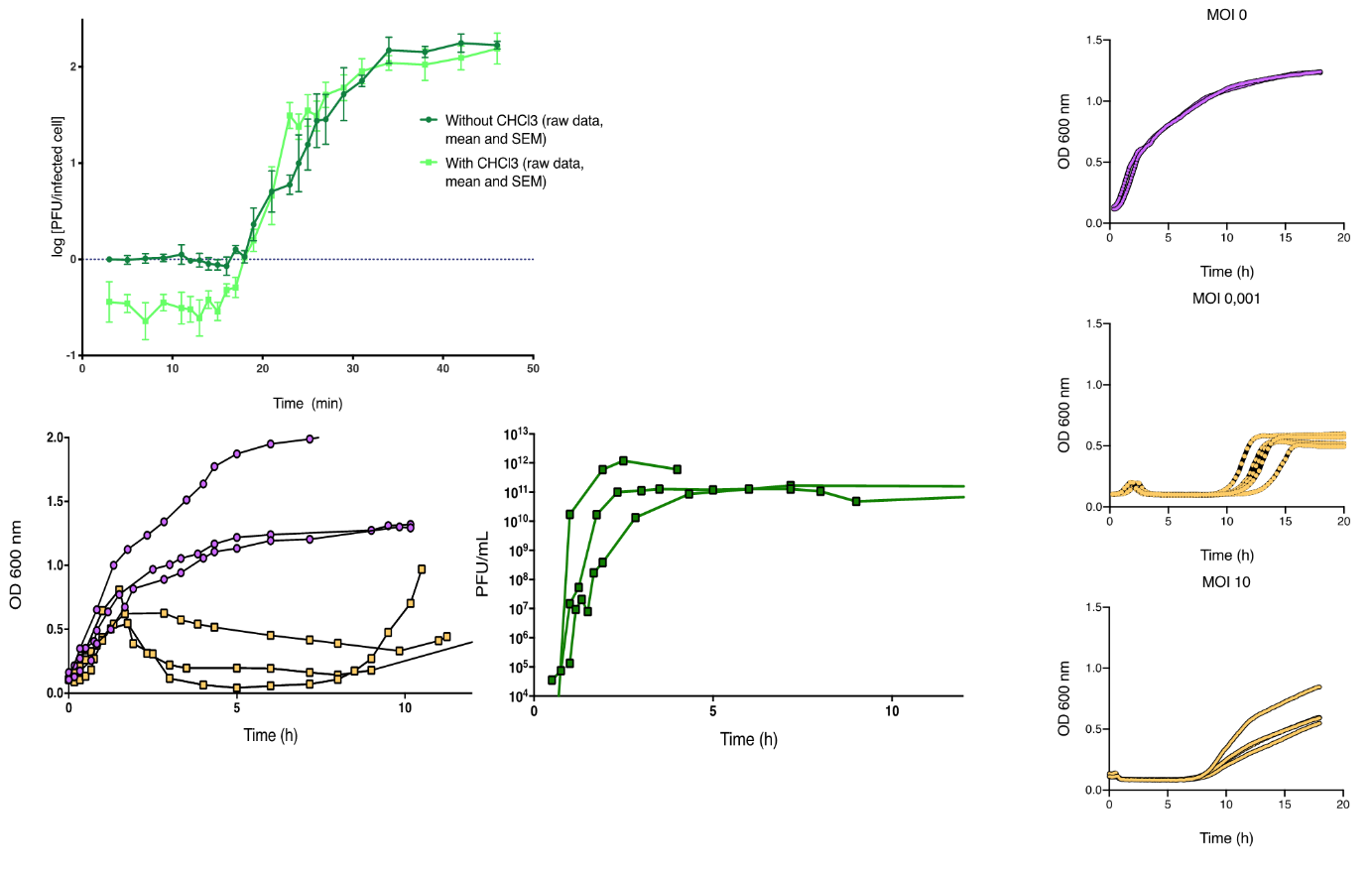

**Supplementary figure S2.** *In vitro* experiments describing the dynamic interactions between strain 536 and phage 536_P1.

The one step growth data (3 replicates) are presented on top left, the dynamic quantification of strain 536 and phage 536_P1 (3 replicates at MOI 0 and at MOI 0.001) are presented on bottom left, and the kinetics of lysis (3 or 4 replicates at MOI 0, MOI 0.0001 and MOI 10) are presented on the right (see Supplementary methods).

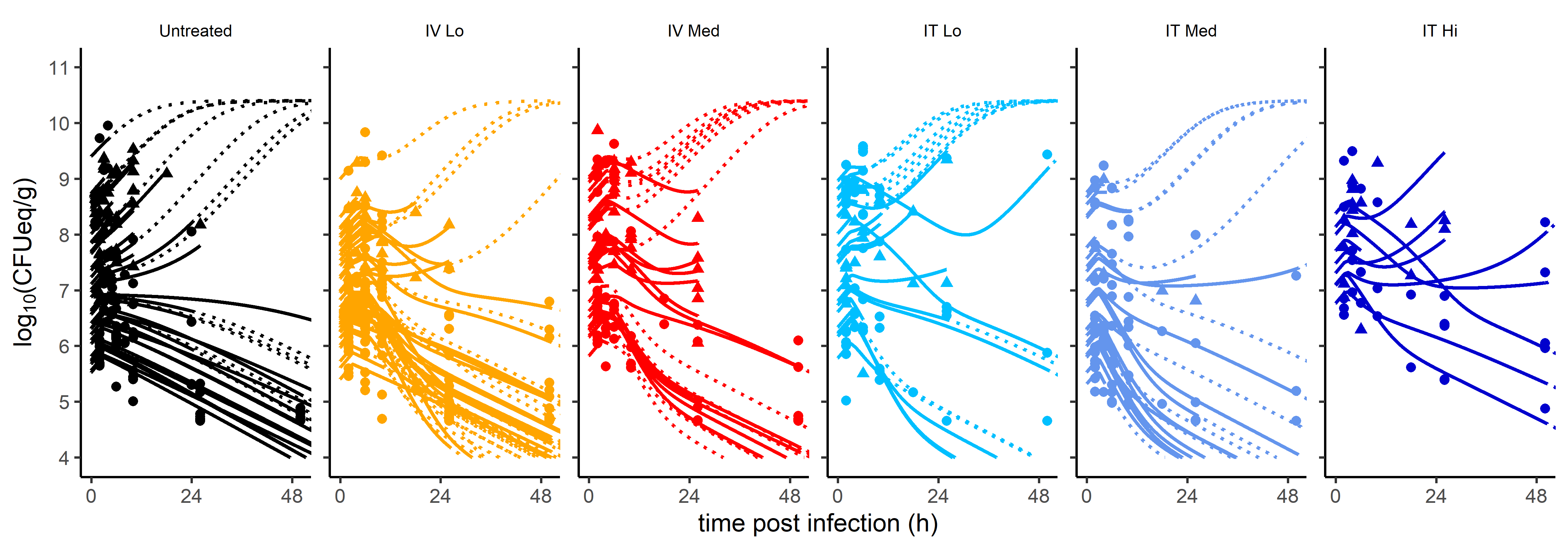

**Supplementary figure S3.** Longitudinal bacterial load data and individual fits from all infected mice (N=183). Solid lines represent individual predictions from the infection time to the last observation time. The dotted lines are individual predictions starting from the sacrifice time point. Interrupted solid lines correspond to animal that die before the scheduled sacrifice time. Circles correspond to live animals while triangles correspond to animals at time of sacrifice after recording the bioluminescence.

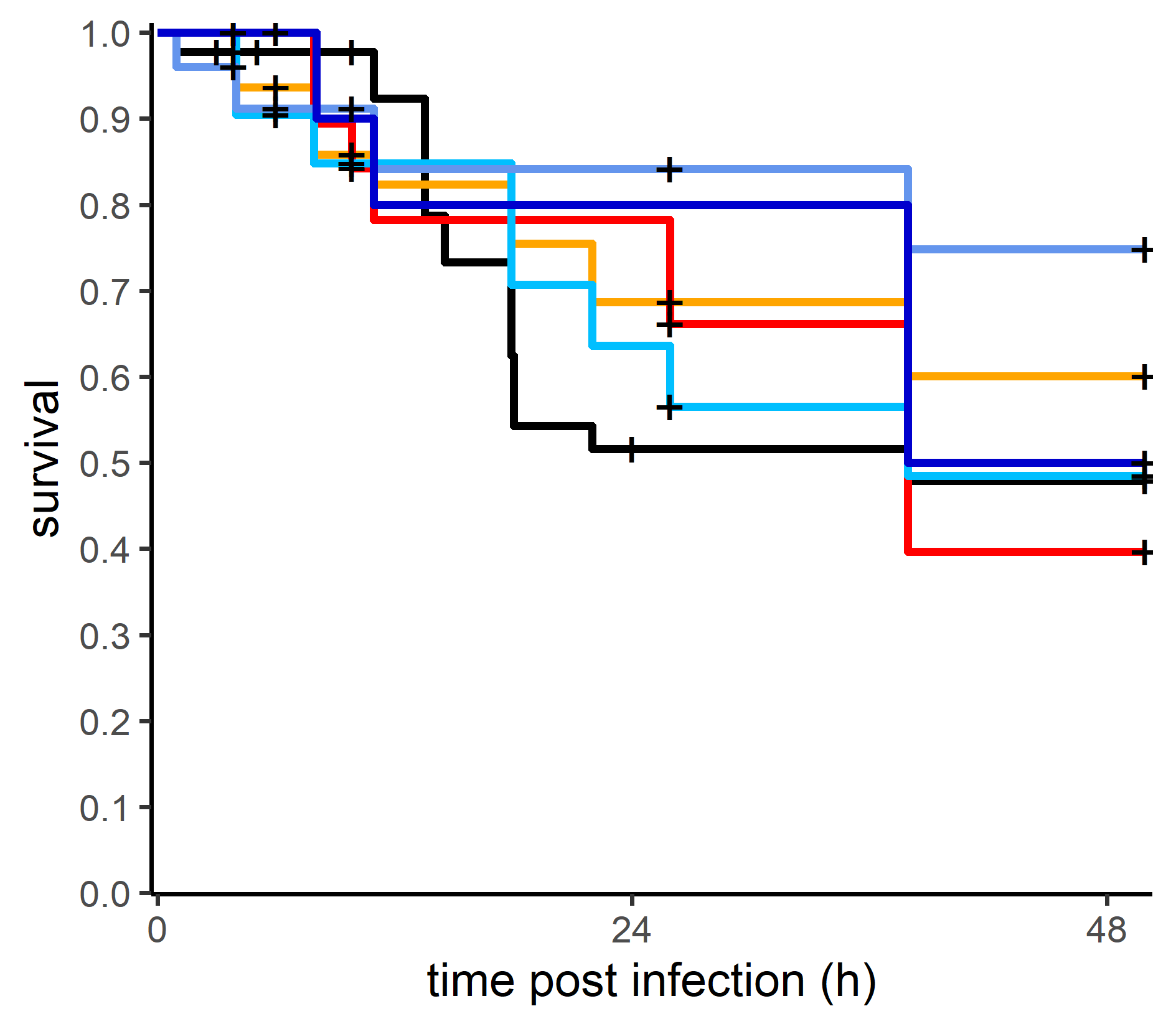

**Supplementary figure S4.** Survival curves over time of *E. coli* strain 536 infected mice and untreated (black, N=45), or treated by phage 536_P1 administered intravenously (IV) at low dose of 5.6 log_10_ PFU (orange, N=50), medium dose of 7.6 log_10_ PFU (red, N=29), intratracheally (IT) at low dose (cyan, N=24), medium dose (blue, N=25) or high dose of 8.6 log_10_ PFU (dark blue, N=10). Crosses represent sacrifice time points.

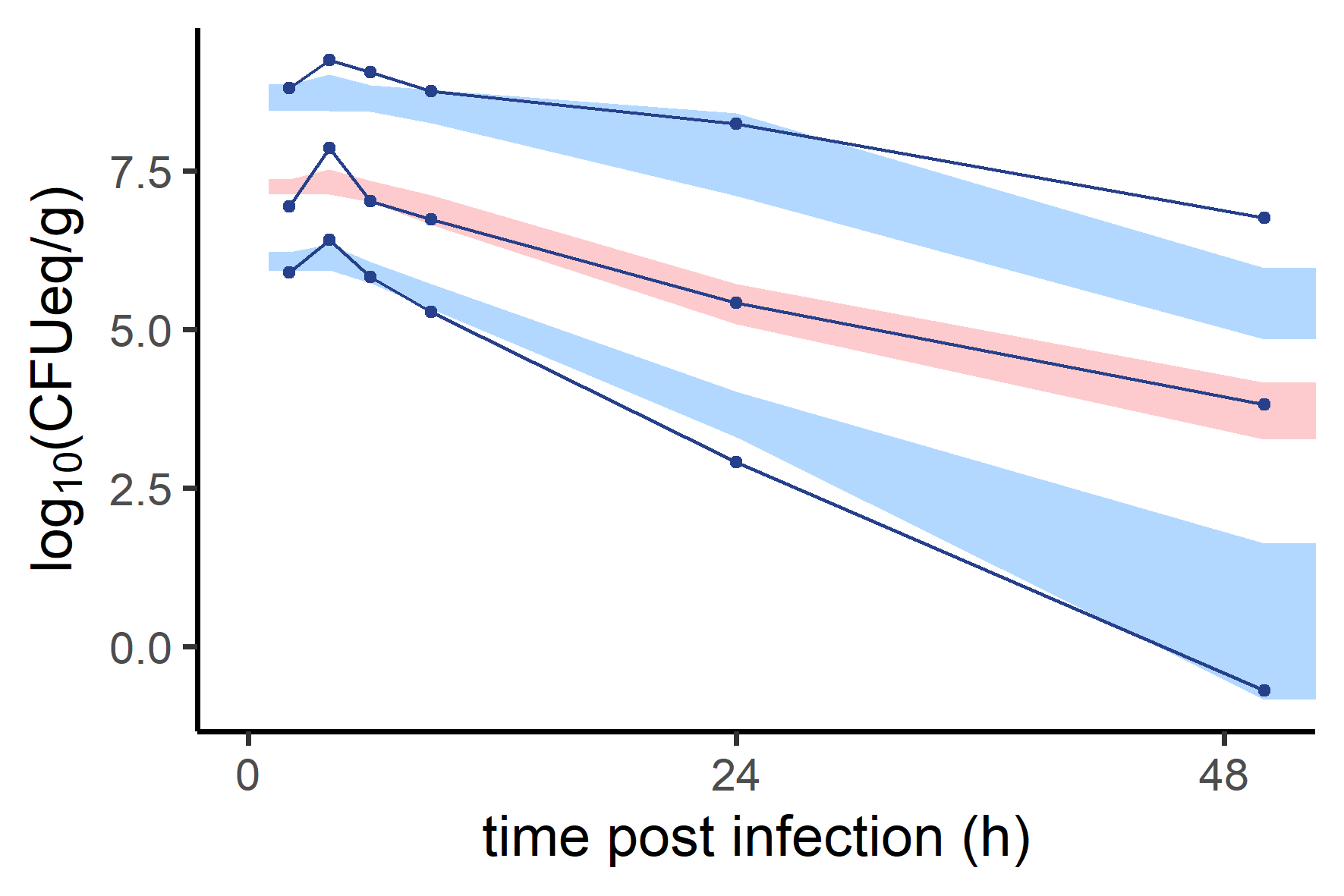

**Supplementary figure S5.** Prediction-corrected visual predictive check plot for the bacterial load, using the final model. The 90% confidence intervals around the 10^th^ (bottom), 50^th^ (middle) and 90^th^ (top) model-predicted percentiles, obtained from 1000 simulated data sets from the model, are represented as blue (10^th^ and 90^th^ percentiles) and pink (50^th^ percentile) areas. The blue dots represent the 10^th^, 50^th^ and 90^th^ percentiles of the observations from the original data set.

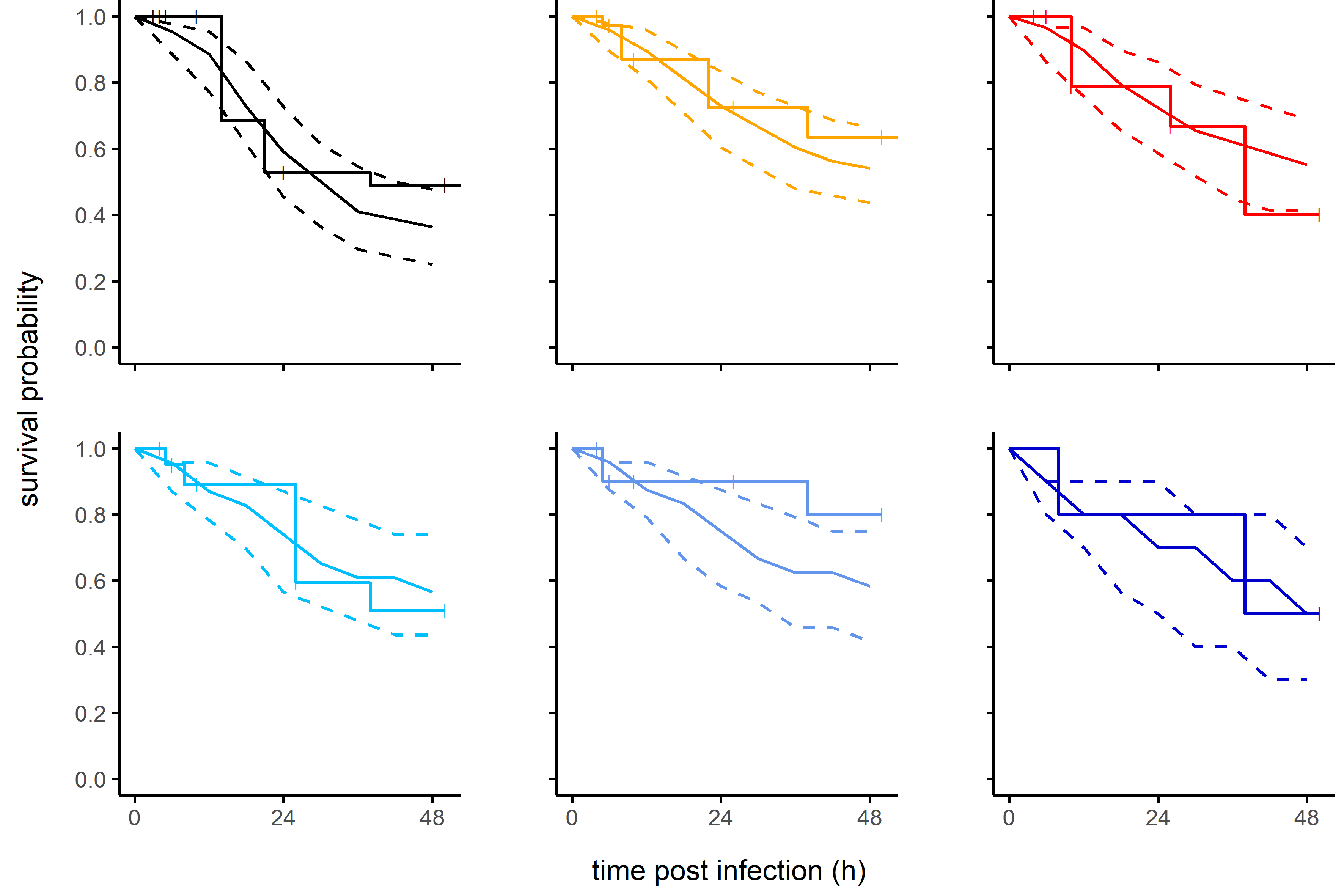

**Supplementary figure S6.** Visual predictive check plot for the survival in each group, using the final model. Untreated mice are in black, phage 536_P1 treated mice (2 h after infection) groups are in colors, with different phage administration scheme: intravenously (IV) at low dose (5.6 log_10_ PFU) in orange or medium dose (7.6 log_10_ PFU) in red, intratracheally (IT) at low dose in cyan, medium dose in blue or high dose (8.6 log_10_ PFU) in dark blue (IT Hi). The dashed lines around the solid line represent the 10^th^ (bottom), 50^th^ (middle) and 90^th^ (top) percentiles of survival obtained from 1000 simulated data sets from the model. The stepped line is the observed survival (Kaplan-Meier curve), with crosses representing censoring.

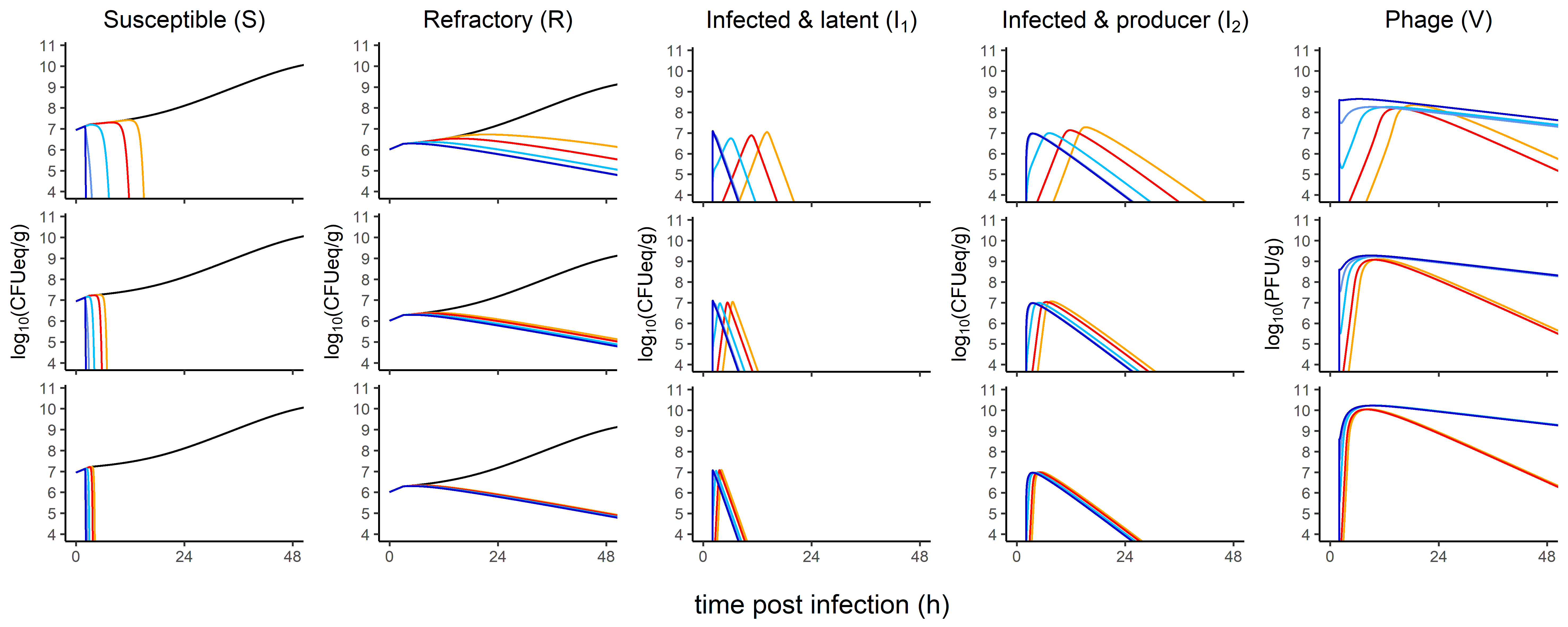

**Supplementary figure S7.** Median profile of model predictions for different burst sizes: 57, 570 (*i.e.* the estimated value) and 5700 PFU/CFU in the top, middle, and bottom line, respectively), with a fixed inoculum of 7 log_10_ CFUeq/g. Each column represents different model compartments: bacteria (S, R, I_1_ and I_2_) and phages in lungs. The colors are indicative of both the route of administration (IT or IV) and the dose of phage received (Lo: 5.6 log_10_ PFU; Med: 7.6 log_10_ PFU; Hi: 7.6 log_10_ PFU): untreated (black), IV-Lo (orange), IV-Med (red), IT-Lo (cyan), IT-med (blue), IT-high (dark blue).

**Supplementary Material**

**Purification of bacteriophages**

Phage 536_P1 was amplified on strain 536 and purified by first, filtration of lysate to 0.1µm followed by tangential ultrafiltration (Vivaflow™, Sartorius, 100kDa), two ultracentrifugations on cesium chloride, dialysed against TN (Tris-HCl pH8, NaCl 150 mM), passed three times through an endotoxin removal column (Endotrap Blue, Hyglos, Germany) and filtered-sterilized at 0.22µm.

***In vivo* experiments**

Intratracheal administration (20µL) was performed in anesthetized mice using elongated tips guided by the glottis visualization with a cold light. The correct position of the tip was checked by the liquid movement inside the tip caused by the ventilatory movement of the animal. The liquid was then expulsed from the tip with a pipette, and 100 µL of air was then pushed to aerosolize the solution to the bronchoalveolar space. Mice were maintained in prone position during at least 5 minutes.

***In vitro* experiments**

**One-Step-Growth experiments**

The one-step growth experiment was performed in triplicate, using Lysogeny Broth (LB, Difco Bacto-Tryptone: 10 g/L, Difco Yeast extract Difco 5 g/L, NaCl 5 g/L). As previously described (Hyman and Abedon, 2009), an exponential-phase culture (9 ml) of strain 536 (optical density at 600 nm 0,25, *ie* 9 log_10_ CFU) was mixed with 1mL of 8 log_10_ PFU of phage 536_P1 (MOI of 0.1). The mixture was placed under constant shaking (100 rpm) at 37°C. Duplicated samples were collected at 2-min intervals during 30 minutes. Six minutes after mixing phages and bacteria, the mixture was 1000 fold diluted in LB medium in order to lower bacterial concentration and avoid any reinfection. Out of the duplicated samples, one was immediately diluted and plated for phage titration, while the second was treated with 10% (vol/vol) chloroform to release intracellular phages in order to determine the eclipse period before phage titration.

**Dynamic quantification of bacteria and phages over time**

We first established the correlation between optical density and CFU for strain 536 by plating on LB agar dilutions of independent aliquots of exponentially growing cells from which OD600 nm was recorded before dilutions. Then, an exponential bacterial culture (50 mL of 1.10^7^ CFU/mL ie a total of 8.7 log_10_ CFU) was mixed with 5.7 log_10_ PFU (ie MOI 1.10^-3^) and incubated under constant shaking (150 rpm, 37°C). Samples from this mixture were withdrawn at specific time points to measure optical density at 600 nm and count the number of phages after serial dilutions and plating on LB agar covered with strain 536. Experiments were performed three times from independent cultures.

**Kinetic of lysis**

To assess the behavior of strain 536 in presence of phage 536_P1 at different MOI, OD600 nm was recorded over time from microtiter plates. In each well, 180 µL of an exponential culture of strain 536 (OD 0.1) was mixed with 20 µL of phage 536_P1 at different concentrations (each replicated six times) to reach MOI of 0.001; 0.01; 0.1; 1 and 10. Controls included wells with phage alone, or bacteria alone or LB alone. The microplates were incubated in a microplate reader (Tecan) set at 37°C during 12h with orbital agitation. Three independent experiments were performed.

***In vitro* data analysis**

Models of *in vitro* data were built using similar methodology used for analyzing *in vivo* PK/PD data: parameters of mixed models were estimated using SAEM algorithm and the model selection was based on BIC. Model of phage/bacteria interactions *in vitro* was built in three steps.

First, one step growth experiments reproduced a single lysis cycle of the phage and therefore allowed to estimate the burst size (*i.e.* the number of virions relased from each bacterium lysed) and the time of a cycle, including an eclipse period. The best fit of these one step growth data was obtained using a multiple transit compartments model. The burst size (570 virions) was estimated with acceptable precision of estimation (relative standard error <30%). The median period for a phage to produce and release new phages was estimated at 35 minutes. Second, bacterial growth was characterized from the above *in vitro* experiments exploiting the condition MOI 0 (without any phage). The bacterial doubling time was estimated at 63 minutes from the model. Third, lysis kinetics experiments (at MOI> 0) were modelled using a predator-prey model with similar structure to *in vivo* model (see equations and table of parameters below). Burst size and bacterial growth were fixed at values respectively obtained at the above first and second steps. The lysis rate (corresponding to rate between infected latent and infected productive bacteria) was fixed from one step growth experiments on 536_P1. The eclipse period was obtained by sensitivity analysis on the transition rate $e$. The proportion of phage-resistant clones at baseline and their relative fitness were characterized from this model.

$$\frac{dS}{dt}=\text{α}\left( 1-\frac{S+I_{1}+I_{2}+R}{{10}^{B_{max}}} \right)\text{S}-\text{β}VS$$

$$\frac{dI_{1}}{dt} =\text{β}VS-eI_{1}$$

$$\frac{dI_{2}}{dt} = eI_{1}- \delta_{V}I_{2}$$

$$\frac{dR}{dt}=w\text{α}\left( 1-\frac{S+I_{1}+I_{2}+R}{{10}^{B_{max}}} \right)\text{R}$$

$$\frac{dV}{dt} = -\text{ }\text{β}VS+\delta_{V}{bI}_{2}$$

Table S1. Parameter estimates of the final *in vitro* model

| **Parameter** | **Name** | **Unit** | **Fixed/Estimated from** | **Fixed effect**  **(rse, %)** | **Sd of random effect (rse, %)** |
| --- | --- | --- | --- | --- | --- |
| Bacterial growth rate | $\text{α}$ | min^-1^ | Coev + Lysis (MOI 0) | 0.011 (-) | 0 (-) |
| Initial number of susceptible | $S_{0}$ | $\log_{10} CFU mL$^-1^ | Coev + Lysis (MOI 0) | 7.9 (-) | 0 (-) |
| Bacterial load plateau | $B_{max}$ | $\log_{10} CFU mL$^-1^ | Coev + Lysis (MOI 0) | 9.9 (-) | 0 (-) |
| Infectivity rate | β | $\log_{10}$ mL CFU^-1^ min^-1^ | Lysis (MOI > 0) | 9.6 (<1) | 0.01 (26) |
| Transition rate | $e$ | min^-1^ | Lysis (MOI > 0) | 0.02 (-) | 0 (-) |
| Lysis rate | $\delta_{V}$ | min^-1^ | Dufour et al., 2017 | 0.5 (-) | 0 (-) |
| Burst size | $b$ | PFU CFU^-1^ | OSG | 570 (-) | 0 (-) |
| Initial number of resistant | $R_{0}$ | $\log_{10} CFU mL$^-1^ | Lysis (MOI > 0) | 6.7 (<1) | 0.05 (10) |
| Relative fitness | $w$ | - | Lysis (MOI > 0) | 0.49 (4) | 0.27 (10) |
| Additive error on bacterial load | $\sigma_{B}$ | $\log_{10} PFU g$^-1^ | Lysis (MOI > 0) | - | 0.1 (1) |

Sd: standard deviation, rse: relative standard error
